# Effect of morphine dependence and withdrawal on operant social interaction in male and female rats

**DOI:** 10.1101/2025.08.28.672892

**Authors:** Emma M. Pilz, Kayla M. Pitts, Jonathan J. Chow

## Abstract

Opioid addiction is linked to decreased social connections. In preclinical models, non-contingent experimenter-administered morphine both decreases unconditioned social interaction and place preference for social reward. We tested if these effects generalize to an operant rat model of social self - administration, in which rats work volitionally for access to a peer. Based on the literature, we also tested if a kappa opioid receptor (KOR) antagonist (LY2456302) and serotonin and dopamine reuptake inhibitors (fluoxetine and GBR12909), would independently reverse the effect of morphine exposure on social self-administration.

We trained rats (n = 66; 32 females) to lever-press for 15-s access to a peer (fixed ratio 1 reinforcement schedule, 45 min, every other day). In Experiments 1-3, we assessed social self-administration during opioid dependence (∼16 h post-injection), and after early (2-to-6 days) and protracted (21-28 days) withdrawal with different morphine exposure regimens (0-to-80 mg/kg, s.c., twice daily; 0-to-80 mg/kg, once daily; or 0-to-40 mg/kg, every-other-day). In Experiment 4, we tested the effects of LY2456302, fluoxetine, and GBR12909 on social self-administration during morphine exposure (every-other-day, 0-to-30 mg/kg).

As in our previous studies, social interaction functioned as an operant reinforcer. Operant social interaction was decreased during morphine exposure (dependence state) but not during early or protracted withdrawal. None of the tested compounds (LY2456302: 5, 10 mg/kg, s.c.; fluoxetine: 1, 3 mg/kg, i.p.; GBR12909: 3, 10 mg/kg, i.p.) reversed this effect.

Opioid dependence, but not withdrawal, decreased operant social interaction in male and female rats. This effect appears independent of KOR, serotonin, or dopamine signaling.

**Highlights:**

- Opioid dependence decreased operant social interaction
- Early and protracted withdrawal had no effect on operant social interaction
- Pharmacological manipulations of Dyn, DA, and 5-HT did not restore social behavior

## Introduction

Opioid addiction is associated with a disruption in social behaviors such as reduced engagement, impaired relationships, and increased isolation (Christie, 2021; Heilig et al., 2016; Inagaki, 2018). These opioid-induced social deficits can persist well beyond the resolution of physical withdrawal symptoms and are hypothesized to increase the likelihood of relapse (Ozdemir et al., 2023; Strang et al., 2020). Despite these observations, insights into the mechanisms underlying the relationship between opioid use disorder and social deficits remain limited.

Studies using mice have shown that decreased social interaction emerge during both experimenter-induced naloxone-precipitated withdrawal and spontaneous withdrawal (for reviews see (Ozdemir et al., 2023; Welsch et al., 2020)). Specifically, studies examining spontaneous withdrawal from opioid agonists have shown that decreased social interaction persist for up to eight weeks after last opioid exposure (Ozdemir et al., 2023; Welsch et al., 2020). In these studies, dependent measures include unconditioned social interactions, such as proximity, physical contacts, and allogrooming when mice are placed together (Becker et al., 2017; Becker et al., 2021; Bravo et al., 2020; Goeldner et al., 2011; Lalanne et al., 2017; Lutz et al., 2014), conditioned place preference for a peer which measures the time spent in a context previously paired with a peer (Pomrenze et al., 2022), and social preference tests where mice choose between a place with a peer versus another without (Becker et al., 2021; Fox et al., 2023; Pomrenze et al., 2022; Zanos et al., 2014).

While previous studies assess the effects of opioid withdrawal on social behavior, less is known about whether opioid dependence and withdrawal affects rewarding volitional social interaction, as assessed in the operant self-administration model, the gold-standard model for evaluating the rewarding effects of drug and non-drug rewards (Wise, 2002, 2004). Operant procedures, where rats are trained to lever-press to access a peer (Angermeier, 1960; Evans et al., 1994), have been used to study social reward. In recent years, these operant methods in rats have re-emerged (Chow et al., 2022; Venniro and Shaham, 2020; Venniro et al., 2018), and, more recently, have been adapted for use CD1 mice (Lee et al., 2025; Ramsey et al., 2022). However, in C57BL/6J mice, the most used background strain for transgenic mice and the strain used in the studies cited above, peer access does not function as an effective operant reinforcer, nor does it elicit conditioned place preference in these mice (Ramsey et al., 2022). This limitation suggests that findings from C57BL/6J mice may not generalize to operant social reward in rats. Therefore, rat models, which reliably show peer-reinforced operant responding, may provide a better framework to study social reward under opioid dependence and withdrawal.

In the present study, we examined the effects of opioid dependence (∼16-18 h post-injection during chronic administration) and withdrawal (up to 4 weeks) on operant social self-administration in male and female rats using different regimens of non-contingent morphine exposure. Additionally, extensive research shows that (1) dynorphin (Dyn) and kappa opioid receptors (KOR) contribute to negative affective states of opioid dependence and withdrawal (Ozdemir et al., 2023; Tejeda and Bonci, 2019), that (2) Dyn suppresses striatal dopamine (DA) and serotonin (5-HT) during opioid dependence and withdrawal, (Goeldner et al., 2011; Spanagel et al., 1992), and that (3) both DA and 5-HT are critical to social behavior (Chow et al., 2024; Kiser et al., 2012; Manduca et al., 2016; Siviy et al., 2011; Vanderschuren et al., 2016). Therefore, based on this literature, we tested if a KOR antagonist (LY2456302), 5-HT reuptake inhibitor (fluoxetine), and DA reuptake inhibitor (GBR12909) could reverse the inhibitory effect of opioid exposure on social self-administration.

## Materials & Methods

### Subjects

For all experiments, we used male and female Sprague-Dawley rats. We used 24 (12 female) rats in Experiment 1, 12 (6 female) rats in Experiment 2, 12 (6 female) rats in Experiment 3, and 18 (8 female) rats in Experiment 4 as experimental rats. We used a total of 66 (32 female) rats as peers. All rats were ∼7-8 weeks PND at arrival (body weight: males, 201-225 g; females, 176-200 g; Charles River). The rats arrived pair-housed and acclimated together for one to two weeks, after which we separated and single-housed the rats. Approximately one week before the start of the experiments, we handled the rats daily. We performed all experiments in accordance with the NIH Guide for the Care and Use of Laboratory (8^th^ edition), under protocols approved by the NIDA IRP Animal Care and Use Committee.

### Apparatus

We conducted the experiments in operant chambers with an attached partner chamber (ENV-00CT-SOCIAL; Med Associates). A guillotine door (ENV-10B2-SOC) and a metal divider with cutouts separated the two chambers, preventing the rats from crossing into the other chamber but allowing for full face-to-face and forepaw contact. We outfitted the front panel of the operant chamber with a recessed pellet receptacle (ENV-200R1M), equipped with a head entry detector (ENV-254-CB), and two retractable levers (ENV-112CM) mounted on either side. A mounted dispenser (ENV-203 M-45) delivered food pellets. Above each lever was a white cue-light (ENV-221 M) and above the recessed pellet receptacle, at the top of the panel, a white houselight. At the very top right corner of the front panel (above the right white cue-light and lever) was a tone generator (ENV-223 AM; 2.9 kHz).

### Drugs

We received morphine sulfate dissolved in sterile saline from the NIDA IRP pharmacy. The NIDA drug supply provided us with LY2456302 (dissolved in 5% DMSO and sterile saline). Dr. Kenner Rice (NIDA IRP) provided us with fluoxetine (dissolved in sterile saline), and GBR12909 (dissolved in sterile saline). The doses of GBR12909, fluoxetine, and LY2456302 are based on previous studies (Bossert et al., 2019; Kelley and Lang, 1989; Sanabria et al., 2008).

### General procedures

On the first day of each experiment, we trained the rats for door shaping followed by magazine shaping. In general, the peer that was paired with each rat was different from arrival and would serve as the peer for rest of the experiment. The door shaping procedure began with a 15-min acclimation phase followed by a phase in which the guillotine door separating the experimental and peer rats was lifted for 30 s, allowing for contact through the cutouts in the metal divider. This repeated for 15 trials every 60 s. Five minutes after the last social interaction trial, we removed the peer rats from the partner chambers and initiated magazine training. The magazine shaping procedure began with a 15 min acclimation phase followed by a phase in which a single pellet would be dropped into the food receptacle every 90 s for a total of 15 pellets. Five minutes after the last pellet, we removed the rats from the operant chambers and returned them to their homecage in the colony.

We then trained rats for social and food self-administration on a fixed-ratio (FR) schedule of reinforcement. Both self-administration sessions functioned similarly such that each session began with the extension of a single lever (counterbalanced across subjects; one lever for social and one lever for food). During each trial, completion of the FR requirement (FR1) resulted in the retraction of the lever, illumination of the cue-light (5.45 s) located above the lever, and delivery of the reward. For social, the guillotine door would open for 15 s; for food, a single 45 mg pellet would be dispensed. To keep the maximal rate of reinforcement equivalent, social trials were separated by a 20 s intertrial interval (ITI), whereas food trials were separated by a 35 s ITI. All sessions lasted 52 minutes. All rats were trained for social, then food self-administration (initiated ∼ 5 to 10 min after the end of social self-administration). Prior to starting food self-administration, the peer-rats were removed f rom the partner chambers and returned to their homecages.

In general, we trained the rats daily during initial self-administration of social and food (detailed below). However, during morphine exposure (detailed below), we trained rats for social and food self - administration every-other-day. This was to account for decreases in social self-administration that occur with daily training, and to dissociate these decreases from the effects of morphine exposure. We injected morphine subcutaneously and altered the injection site between days. All peer rats remained morphine naïve throughout all experiments. We weighed the rats daily before the start of the self-administration sessions.

#### Experiment 1 – Effect of repeated rounds of morphine exposure

In Experiment 1, we trained the rats for four consecutive daily sessions with either a same- or opposite-sex peer during social self-administration (Fig 1A). We then gave the rats 1-2 d breaks and then trained them with the other peer-sex. We trained rats on this pattern for 12 to 16 sessions. At the end of initial self-administration training, rats remained paired with whichever peer sex they finished training with.

**Figure 1.**
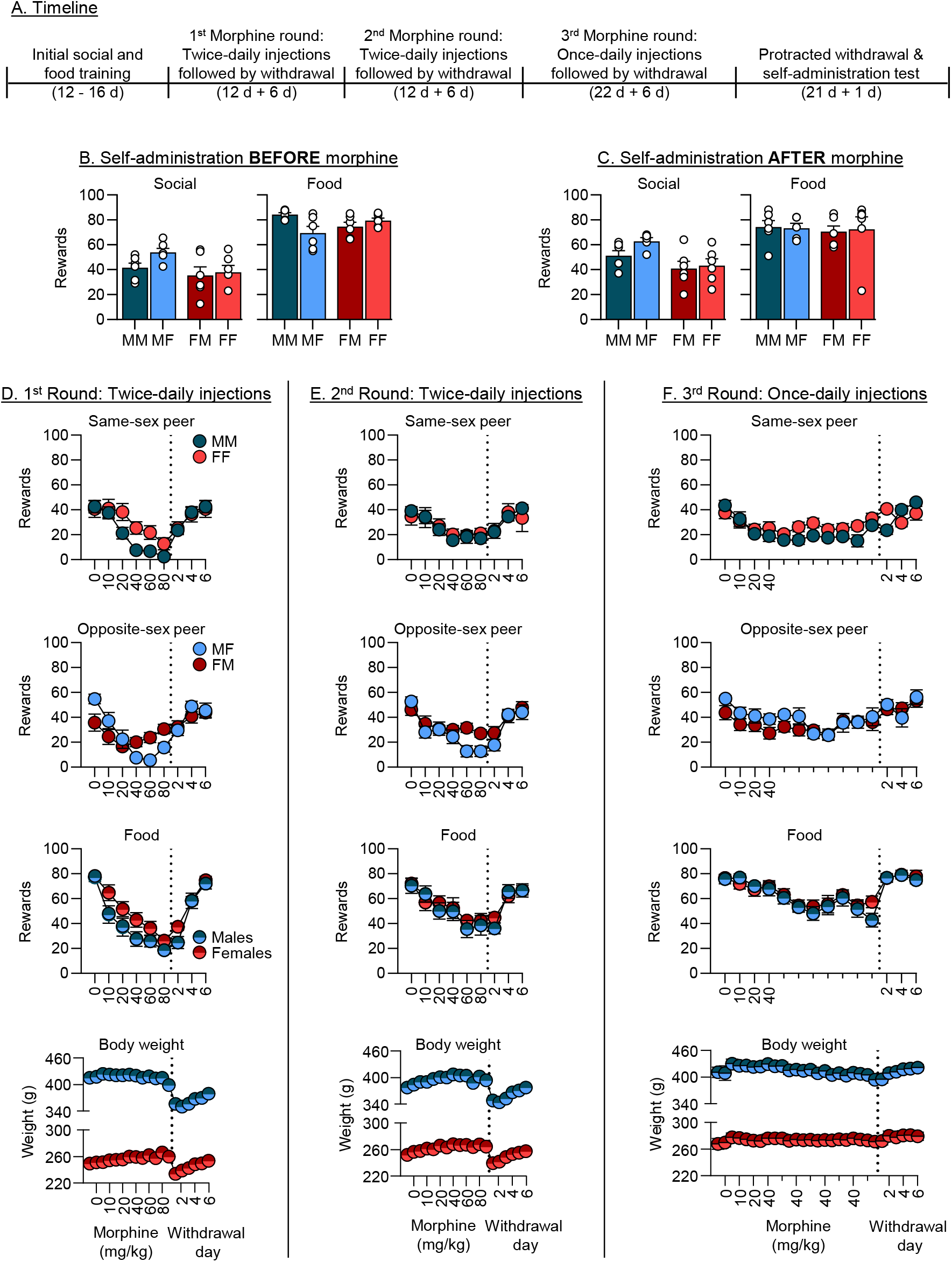
Effect of repeated rounds of morphine exposure. (**A**) Representative timeline of Experiment 1. Number of social and food rewards earned during self-administration (**B**) before and (**C**) after repeated rounds of morphine injections. The circles representative individual data points. Note: The first letter denotes sex of the experimental rat (i.e., M = male, F = female) and the second letter denotes the peer-sex. (**D, E, & F**) Self-administration across the three rounds of morphine injections for a same-sex peer (topmost panels), opposite sex peer (second from top panels), food (second from bottom panels), and weight (bottom most panels). All datapoints left of the dotted line represent self-administration during morphine dependence, while datapoints on the right of the line represent self-administration during early withdrawal. Minor ticks represent continuation of last major tick. Notes: Withdrawal sessions 2, 4, 6 are approximate days since last injection. Scale for males and females are different (increments of 60 in males vs. 40 in females).

For Experiment 1, we exposed the rats to three rounds of morphine exposure. These rounds refer to increasing morphine doses over days and then ceasing morphine injections and allowing the rats to go through spontaneous withdrawal. We based our initial morphine exposure regimens on previous literature (Anraku et al., 2001; Linseman, 1977). For the first two rounds, we used an injection schedule where rats were injected twice daily; rats received each dose for two days (0, 10, 20, 40, and 80 mg/kg morphine; over 12 d). Social and food self-administration training occurred on the second day of injections at each dose. We injected rats around the same time which coincides with the end of self-administration training in the morning and then again ∼7 h later (injections at ∼10 am and ∼5 pm); rats were trained for self-administration ∼16 h since last injection. After the morphine injections, we tested the rats during early withdrawal (over 6 d, every-other day); the first self-administration session during the withdrawal period occurred ∼40 h after last morphine injection. For the third round, we tested whether once-daily morphine injections could maintain ∼50% suppression of operant responding for social interaction using once-daily injections. We escalated (0, 10, and 20 mg/kg) and maintained the rats at 40 mg/kg morphine for 22 d (injections at ∼5 pm; social self-administration starts ∼16 h since last injection) and then tested the rats during early withdrawal over 6 d (every-other-day) with the first self-administration session during the withdrawal period being ∼40 h since last injection. Finally, we tested the rats 21 d (28 d since last injection) later for social and food self-administration to determine if protracted withdrawal can result in long-term social deficits.

We removed one male rat paired with a female-peer from this experiment due to unexpected death.

#### Experiment 2 – Effect of daily experimenter-administered morphine exposure

In Experiment 2, we wanted to determine if we could replicate a sustained ∼50% suppression in social interaction, ideally replicating the third round of once-daily morphine injections in Experiment 1 (see Fig 2A for timeline). We paired rats with a same-sex peer for social interaction (and for the rest of the subsequent experiments); this was done because we observed no differences in peer-sex during the once-daily injection schedule in Experiment 1. We trained the rats for social and food for four daily consecutive sessions with a 1-2 d break in between bouts of daily training; we trained rats for 12 sessions total.

**Figure 2.**
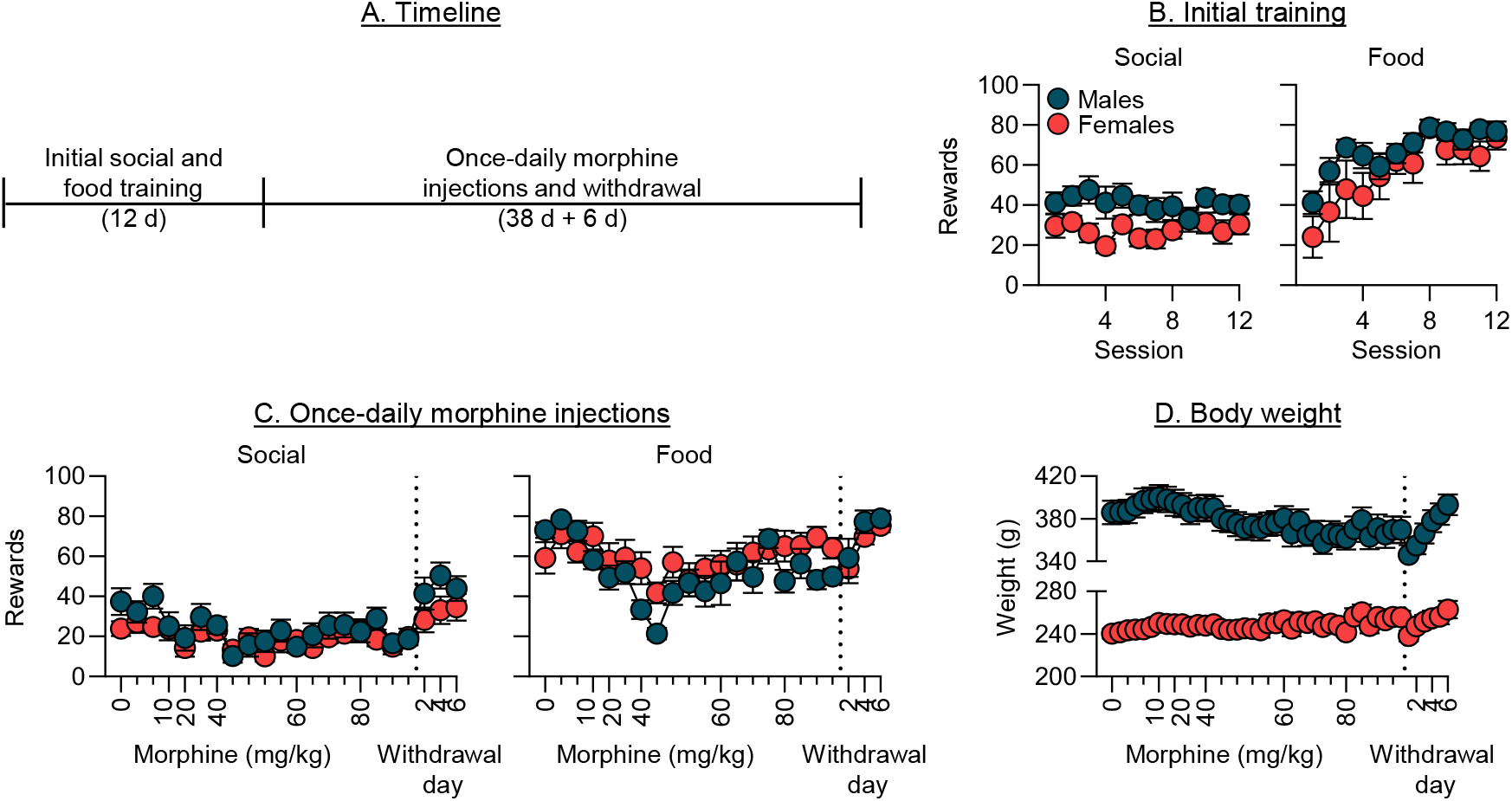
Effect of once-daily morphine exposure. (**A**) Representative timeline of Experiment 2. (**B**) Initial social and food self-administration before morphine injections. (**C**) Social and food self-administration during morphine injections (left of dotted line) and during early withdrawal (right of dotted line). (**D**) Body weight throughout the experiment. Minor ticks represent continuation of last major tick. Withdrawal sessions 2, 4, 6 are approximate days since last injection.

In Experiment 2, we injected rats with saline (0 mg/kg) for 6 d, 10 mg/kg morphine for 2 d, 20 mg/kg morphine for 4 d, 40 mg/kg morphine for 10 d, 60 mg/kg morphine for 8 d, and 80 mg/kg morphine for 8 d while training the rats for social and food self-administration (injections at ∼3 pm; self-administration starts ∼18 h after last injection). After, we stopped all morphine injections and continued to train rats for social and food self-administration over a 6-d period (every-other-day) with the first self-administration session during the withdrawal phase being ∼32 h since last injection. We increased the dose of morphine during the 38-d period based on positive trends in the number of social interactions earned during a given dose of morphine.

#### Experiment 3 – Effect of every-other-day morphine exposure

Given the observed tolerance with daily morphine injections in Experiment 3, we wanted to determine if using an every-other-day injection schedule could maintain ∼50% suppression of operant social interaction (Fig 3A). We initially trained rats for social and food self-administration for four consecutive daily sessions with a 2-d break before training them for five consecutive daily sessions.

**Figure 3.**
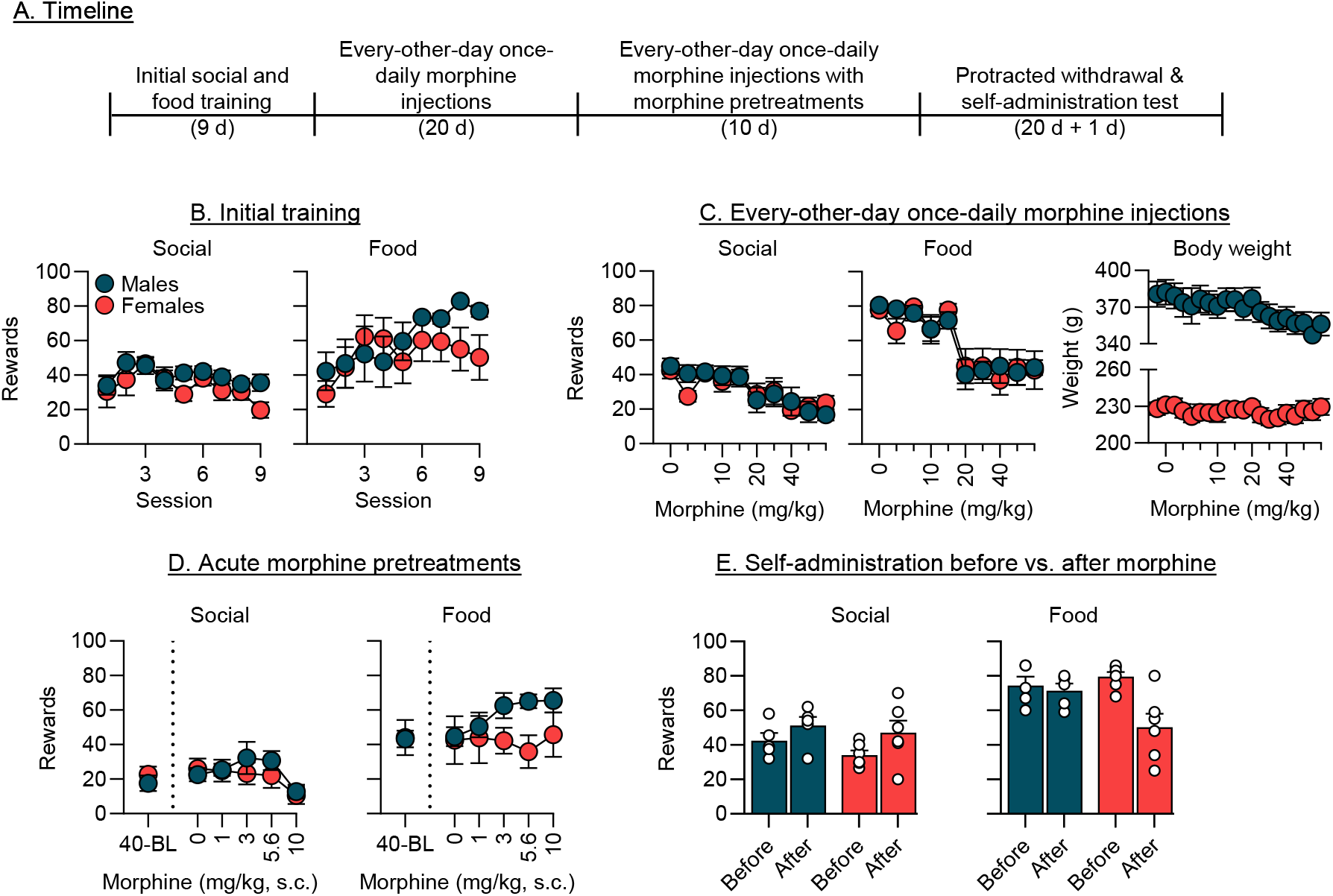
Effect of every-other-day morphine exposure. (**A**) Representative timeline of Experiment 3. (**B**) Initial social and food self-administration before morphine injections. (**C**) Social and food self-administration during morphine injections with corresponding weights. Minor ticks represent continuation of last major tick. (**D**) Effect of acute morphine pretreatments prior to social and food self-administration. 40-BL (baseline) represents the average of the last 2-d of self-administration at 40 mg/kg morphine prior to pretreatments. (**E**) Number of social and food rewards earned during self-administration (**B**) before and (**C**) after every-other-day morphine injections.

Next, we injected the rats every-other-day with (0 mg/kg) over a 6-d period, 10 mg/kg morphine over a 4-d period, 20 mg/kg morphine over a 4-d period, and 40 mg/kg morphine over a 6-d period.

Specifically, we only injected rats the day before social and food self-administration (injections at ∼ 3 pm; self-administration starts ∼18 h after last injection; ∼48 h between injections), such that on the days that we trained the rats they received no injections. Next, while maintaining rats on 40 mg/kg morphine on an every-other-day schedule, we pretreated rats with morphine (0, 1, 3, 5.6, and 10 mg/kg s.c.; pseudo-Latin square design) 30 minutes prior to the start of social self-administration. After the last morphine pretreatment day, we ceased all injections and then retested the rats for social and food self - administration 21 d later.

We removed one male rat from this experiment due to unexpected death.

#### Experiment 4 – Effect of GBR12909, fluoxetine, and LY2456302 on morphine-induced suppression of social and food self-administration

In Experiment 4, we tested if GBR12909, fluoxetine, and LY2456302 would reverse morphine-induced suppression of social self-administration (Fig 4A). We trained rats for social and food self-administration for four consecutive daily sessions with a 1-d break before training them for five consecutive daily sessions. Next, to assess the effects of the pharmacological agents on social and food self-administration prior to morphine exposure, we continued to run the rats for five consecutive daily sessions. During this phase, in half of the rats we pretreated them with fluoxetine (0, 1, and 3 mg/kg, i.p.) and the other half with GBR12909 (0, 3, 10 mg/kg, i.p.) 30 min prior to the start of social self - administration. We tested doses in ascending order and gave the rats a three-session washout period. We then tested the rats previously treated with GBR12909 with LY2456302 (0, 5, and 10 mg/kg, s.c.; 2-h pretreatment); we continued to the run the rats pretreated with fluoxetine. We only tested half the rats initially as we did not have enough LY2456302 to test all rats.

**Figure 4.**
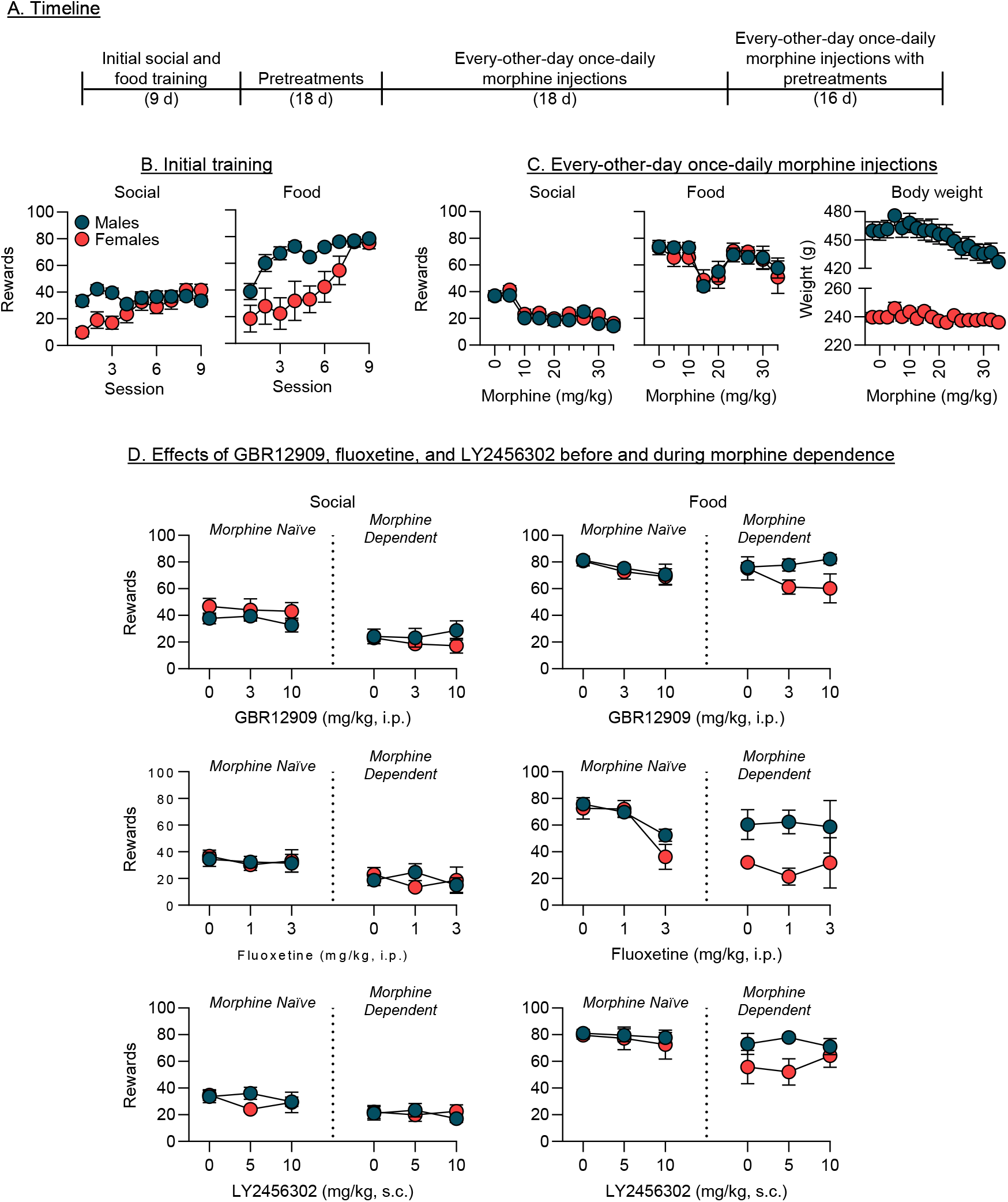
Effect of GBR12909, fluoxetine, and LY2456302 on morphine-induced suppression of social and food self-administration. (**A**) Representative timeline of Experiment 4. (**B**) Initial social and food self-administration before morphine injections. (**C**) Social and food self-administration during morphine injections with corresponding weights. Note: scale for males and females are different (increments of 30 in males vs. 20 in females). Minor ticks represent continuation of last major tick. (**D**) Effect of GBR12909 (top), fluoxetine (middle), and LY245302 (bottom) on social and food self-administration prior to (left of dotted line) and during morphine exposure (right of dotted line).

Next, we started morphine injections in the rats using an every-other-day injection schedule (injections at ∼ 3 pm; self-administration starts ∼18 h after last injection; ∼48 h between injections) with (0 mg/kg) over a 4-d period, 10 mg/kg morphine over a 4-d period, 20 mg/kg morphine over a 6-d period, and 30 mg/kg morphine over a 4-d period while training the rats for social and food self-administration on the off-days. We stopped at 30 mg/kg morphine for this experiment because we had 2 rats (1 male, 1 female) that unexpectedly died after 10 mg/kg morphine injections and another rat (male) that died after 20 mg/kg morphine injections. We maintained rats on 30 mg/kg morphine. During this phase, we pretreated the same rats that received fluoxetine and GBR12909, respectively, at the same drug and doses previously used but in pseudo-random order. We then tested pretreatments of LY2456302 in all rats in pseudo random order of dose.

## Data analysis

We analyzed the data using linear mixed-effects (LME) model in JMP (version 16) where we set α To 0.05. In all behavioral analyses, we used subjects (the rats) as a random factor. Importantly, because morphine doses increased across self-administration sessions, and doses were typically repeated, we analyzed the morphine exposure data using the number of training sessions (continuous) but graphically displayed the corresponding dose; we did the same for early withdrawal. We followed up on significant interactions, when allowed, using Tukey’s HSD post-hoc test. We describe the factors for each statistical analysis in the “Results” section. We only report significant effects critical for data interpretation and indicate results of post-hoc analyses in the figures; all statistics can be found in Table 1.

## Results

### Experiment 1 – Effect of repeated rounds of morphine exposure

We observed no long-term decreases in the number of social and food rewards earned prior to (Fig 1B) vs. 3-weeks after repeated rounds of morphine exposure (Fig 1C). We analyzed the data by comparing the number of social rewards earned prior to morphine (average of last 2-d of training) vs. number of rewards earned during the self-administration test with sex (nominal; male vs. female) and peer-sex (nominal; male vs. female) as between-subject factors and morphine exposure (nominal; before vs. after) as a within-subjects factor. For food rewards, we used the same model but excluded peer-sex as a factor as there was no peer present; we did this for all subsequent analyses for food. This analysis showed a significant main-effect of sex [F(1,20.15) = 7.5, *p* = 0.012] and morphine exposure [F(1,19.33) = 21.4, *p* = 0.0002] for social self-administration, indicating that male rats earned more social rewards than females, and that, in general, there was an increase in social rewards earned after repeated rounds of morphine exposure. There were no significant differences for food self-administration.

The twice-daily and once-daily (maintained up to 40 mg/kg) rounds of morphine injections significantly suppressed social and food self-administration, and cessation of morphine resulted in a rebound in self-administration during early withdrawal (Fig 1D-F). We first analyzed the number of social and food rewards earned during the morphine injections for each round individually with sex and peer-sex (excluded for food) as between-subjects factors, and session as a within-subjects factor. These analyses showed a consistent sex x session interaction across all rounds of morphine injections for social rewards [Round 1: F(1,20.04) = 18.4, *p* = 0.0004; Round 2: F(1,19) = 10.1, *p* < 0.005; Round 3: F(1,19) = 10.8, *p* = 0.004], indicating males decreased responding for a peer to a greater degree than females during opioid dependence. The analysis also showed a main effect of session across all rounds of morphine for food rewards [Round 1: F(1,21.72) = 225.4, *p* < 0.0001; Round 2: F(1,21) = 32.4, *p* < 0.0001; Round 3: F(1,21) = 77.2, *p* < 0.0001], indicating food self-administration was also suppressed.

We then analyzed the number of social and food rewards earned during early withdrawal using the same respective models. These analyses showed that the number of social rewards [Round 1: F(1,19) = 32.9, *p* < 0.0001; Round 2 F(1,19) = 46.2, *p* < 0.0001; Round 3: F(1,19) = 11.4, *p* = 0.0032] and food rewards [Round 1: F(1,21) = 154.6, *p* < 0.0001; Round 2 F(1,21) = 39.5, *p* < 0.0001] significantly increased. It should be noted that no significant differences were observed for food self-administration during the third round as it immediately rebounded back to baseline levels during the first session.

### Experiment 2 – Effect of once-daily morphine exposure

Rats self-administered social and food rewards (Fig 2B). We analyzed the data with sex as a between-subjects factor and session as a within-subjects factor. The analysis showed a main effect of sex for both social [F(1,10.01) = 6.2, *p* = 0.031] and food [F(1,10) = 5.8, *p* = 0.037], indicating that males earned more rewards for females, and a main effect for session during food self-administration [F(1,10) = 13.4, *p* = 0.004], indicating rats increased their food intake over time.

Once-daily injections of morphine resulted in rats showing signs of tolerance (Fig 2C). We analyzed the data for morphine injections using the same model and saw no significant main effects. In addition, we observed trends for increases in social and food rewards at the repeated 40 and 60 mg/kg doses for morphine injections (see Table 1).

### Experiment 3 – Effect of every-other-day morphine exposure

Rats learned to self-administer social and food rewards (Fig 3B). We analyzed the data with sex as a between-subjects factor and session as a within-subjects factor. The analysis showed a main effect of session for both social [F(1,10) = 6.2, *p* = 0.032] and food [F(1,10) = 10.2, *p* = 0.01].

Every-other-day injections of morphine significantly suppressed social and food self -administration (Fig 3C). We analyzed the number of social and food rewards earned during the morphine injections with sex as a between-subjects factor and session as a within-subjects factor. The analysis showed a main effect of session for social [F(1,9.5) = 43.4, *p* < 0.001] and food [F(1,10.4) = 37.2, *p* = 0.001].

Acute morphine pretreatments decreased responding for social interaction but not food (Fig 3D). We analyzed the data with sex as a between-subjects factor and acute morphine dose (continuous; 0, 1, 3, 5.6, and 10) and the analysis showed a main effect of dose [F(1,9) = 13.5, *p* = 0.005] on social interaction; this effect was driven by the 10 mg/kg pretreatment of morphine suppressing responding. There were also no significant differences between a baseline (last 2-d training before acute morphine pretreatments at every-other-day morphine injections) vs. the acute morphine dose at 0 mg/kg (saline).

We observed no long-term decreases in the number of social rewards earned (Fig 3E) prior to vs. ∼3 weeks after last morphine exposure. We analyzed the data with sex as a between-subjects factor and morphine exposure as a within-subjects factor and the analysis showed no significant differences. When we analyzed the data for food, there was a significant effect of morphine exposure [F(1,9) = 6.8, *p* = 0.029]; post hoc analysis showed that this effect was driven by a decrease in food responding in the female rats.

### Experiment 4 – Effect of GBR12909, fluoxetine, and LY2456302 on morphine-induced suppression of social and food self-administration

Rats learned to self-administer social and food rewards (Fig 4B). We analyzed the data with sex as a between-subjects factor and session as a within-subjects factor. The analysis showed a main effect of sex [F(1,16) = 4.7, *p* = 0.045] and session [F(1,16) = 12.7, *p* = 0.0026] and a sex x session interaction [F(1,10) = 16.4, *p* = 0.0009] for social interaction, indicating that male rats earned more social rewards for females, while females increased the number of social interactions per session over time. The analysis also showed a main effect for sex [F(1,16) = 12.2, *p* = 0.003] and session [F(1,16) = 37.9, *p* < 0.0001] for food, indicating that males earned more food rewards than females, and the rats increased food self - administration over training.

Every-other-day injections of morphine decreased social and food self-administration. Using the same model above, the analysis showed a main effect of session for social [F(1,15.47) = 106.7, *p* < 0.0001] and food [F(1,15.13) = 5.3, *p* = 0.036] self-administration.

Pretreatments with GBR12909, fluoxetine, and LY2456302 had no effect on social self-administration and little effect on food self-administration when rats were morphine naïve (Fig 4D, left side). We analyzed the data with sex as a between-subjects factor and dose as a within-subjects factor. The analysis showed no significant effects for any of the compounds on a social self-administration. The analysis did show a main effect for dose for GBR12909 [F(1,7) = 5.6, *p* = 0.05] and fluoxetine [F(1,7) = 29.1, *p* = 0.001], indicating that these compounds dose-dependently decreased food self-administration.

Pretreatments with GBR12909, fluoxetine, and LY2456302 had no effect on social or food self-administration when rats were morphine dependent (Fig 4D, right side). We used the same statistical model above and the analysis showed no significant differences for either social or food self - administration.

## Discussion

We examined the effects of opioid dependence and withdrawal on operant social interaction. We included food self-administration as a relative comparison to evaluate how opioid dependence and withdrawal generally affect motivated behavior. Across the different experimenter-administered morphine regimens, we consistently observed a suppression of responding for both rewards during opioid dependence (i.e., when the rats were receiving morphine). When morphine injections were discontinued, the rats returned to baseline levels of self-administration during early withdrawal (within ∼6 days after the last morphine injection). Furthermore, when assessed during protracted withdrawal (∼3 to 4 weeks after the last morphine injection), we did not observe any notable decreases in social self-administration. Finally, the suppression of social interaction and food self-administration during opioid dependence appeared to be independent of DA, 5-HT, and Dyn neurotransmission.

### Effect of opioid dependence and withdrawal on operant social interaction

We combined a procedure for operant social interaction (Chow et al., 2022; Venniro and Shaham, 2020; Venniro et al., 2018) with different regimens of experimenter-administered morphine known to induce opioid dependence and withdrawal (Anraku et al., 2001; Becker et al., 2010; Linseman, 1977). We initially included peer sex as a variable (Experiment 1) to examine its effects on operant responding for social interaction during opioid dependence and withdrawal. In agreement with our previous findings (Chow et al., 2024), we observed that the male rats lever-pressed more for a female peer than for a male peer when trained with both sexes. However, during opioid dependence and withdrawal, our results did not indicate an effect of peer sex on the number of social rewards earned. Therefore, we used same-sex peers for the remaining experiments. Independent replications and additional research are nevertheless warranted to determine whether peer sex influences other behavioral measures of operant social motivation (e.g., progressive ratio, economic demand, choice between social and food reward) during opioid dependence and withdrawal.

Many studies using chronic experimenter-administered morphine have examined behavioral and neural changes during either early or late (protracted) opioid withdrawal (Ozdemir et al., 2023; Welsch et al., 2020). This raises a caveat for our study regarding the opioid withdrawal state of the rats, particularly during morphine exposure (i.e., opioid dependence). Specifically, in our experiments, we did not observe drastic weight loss between morphine injection days, diarrhea in the homecages or operant chambers, or signs of irritation when the rats were handled—common indicators of opioid withdrawal (Blasig et al., 1973). Thus, we presume the rats were not undergoing bouts of spontaneous withdrawal between injections.

Our data differ from prior research (Martin et al., 1963; Ozdemir et al., 2023; Parker and Radow, 1974) showing that opioid regimens like those we used can induce spontaneous withdrawal symptoms as early as 8 h after the last morphine injection. However, we did observe a large decrease (∼20 to 35 grams) in body weight in the ∼40 h following cessation of morphine exposure (see Figs 1 and 2), a reliable symptom of spontaneous opioid withdrawal (Blasig et al., 1973; Shaham et al., 1996). Of note, under the every-other-day injections (∼48 h between injections), we did not observe the same drop in weight seen under every-day injections going into withdrawal (∼40 h since last injection). Instead, we observed a slow decrease in weight in males, but not females, over the course of the experiment; like the third round of Experiment 1 and Experiment 2. Based on the lack of large weight loss, steady weight in females, and lack of other identifiable symptoms (i.e., diarrhea and irritability), we presume the rats under the every-other-day injections were likely not undergoing spontaneous withdrawal between injections. Nonetheless, future studies using every-other-day regimens should examine the timeline in which major weight loss is observable or if other markers of withdrawal are observed.

In Experiment 3, we tested whether pretreatment with low doses of morphine could restore social and food responding during morphine exposure. We reasoned that such doses might alleviate some psychological or physiological withdrawal symptoms, thereby restoring normal motivation for social self - administration. However, under our experimental conditions, acute morphine injections had no effect. The reasons for these negative results are unknown.

The current literature on spontaneous withdrawal following chronic morphine exposure in mice consistently shows long-term decreases in social behavior under various social procedures (see Introduction; Ozdemir et al., 2023; Welsch et al., 2020). In contrast, the limited number of rat studies examining social interaction after chronic morphine exposure do not show reliable decreases in social interaction after cessation of morphine (Blatchford et al., 2005; Grasing et al., 1996). For example, in Grasing et al (1996), rats were implanted with morphine-filled osmotic minipumps (44 mg/kg/d) for 7 days. Following minipump removal, social interaction was assessed as a proxy for anxiety by placing rats together. Across 3 days of post-morphine testing, there were no decreases in social interaction relative to saline controls. Similarly, in Blatchford et al (2005), rats received daily morphine injections (10 mg/kg, s.c.) for 10 days followed by 7 days of withdrawal. Social interaction, again assessed as a proxy for anxiety, did not decrease during the withdrawal period unless rats experienced an additional stress manipulation. Although these studies did not extend testing to 8 weeks, it seems unlikely that decreases in social interaction would emerge over time. These prior results parallel our present findings, in which operant responding for social behavior was not decreased after cessation of morphine exposure.

Taken together, in contrast to findings from mouse studies but consistent with several rat studies, we found little evidence that either early or protracted opioid withdrawal produces social deficits, as measured by our operant social self-administration procedure. An important question for future research is whether opioid dependence or withdrawal-induced social deficits would emerge in CD1 mice trained in an operant social self-administration procedure (Lee et al., 2025; Ramsey et al., 2022) modeled after the rat procedure used in the present study (Venniro and Shaham, 2020).

### Effect of pharmacological manipulation of DA, 5-HT, and Dyn transmission on social self-administration

One of our initial goals was to examine whether manipulations of DA, 5-HT, and Dyn transmission could restore operant social interaction during opioid withdrawal. However, given the lack of sustained decreases in operant responding for both social and food rewards after cessation of morphine exposure, we instead tested the pharmacological manipulations during opioid dependence, where we observed a reliable decrease in social self-administration. We did not observe any effects of systemic injections of GBR12909, fluoxetine, or LY245302 on morphine exposure-induced decreases in social self-administration.

Despite evidence implicating DA, 5-HT, and Dyn in opioid dependence and withdrawal (Ozdemir et al., 2023; Tejeda and Bonci, 2019) as well as motivated social behavior (Chow et al., 2024; Kiser et al., 2012; Manduca et al., 2016; Siviy et al., 2011; Vanderschuren et al., 2016), our systemic manipulations of each individual neuromodulator had no effect. These negative results suggest that independent systemic manipulations of DA, 5-HT, and Dyn is insufficient to reverse morphine-induced social deficits, raising the question of whether systemic injections of different drug combinations could reverse the effects of opioid dependence on social (and/or food) self-administration.

### Translational considerations – social deficits

In humans, the association between opioid addiction and disruptions in social connection does not exist in a vacuum. It is shaped by personal relationships and influenced by environmental factors such as culture and societal norms, which can precede and exacerbate opioid use (Nicholson Jr, 2023; Shah et al., 2017). Opioid addiction often disrupts relationships with family or long-term support networks while simultaneously fostering new connections within communities of shared drug use (Valentine, 2009). In such cases, cessation of opioid use, aside from physical withdrawal, may create situations where the absence of prior social connections exacerbates negative affective states associated with isolation and loneliness (McIntosh and McKeganey, 2000). While preclinical models provide valuable insight into the neurobiological mechanisms underlying opioid-induced social deficits, they fall short of capturing the complex interactions between opioid addiction and social behavior in humans.

### Conclusions

Our results show that, contrary to previous findings in mice, under our experimental conditions, rats do not exhibit deficits in operant social interaction during opioid withdrawal. Instead, rats show decreased operant responding during opioid exposure and dependence. Additionally, our initial pharmacological investigation suggests that contrary to our hypothesis and the literature, DA, 5-HT, and Dyn transmission do not play a significant role in mediating opioid-induced decreases in social self-administration.

## Acknowledgements

We thank Yavin Shaham for comments on earlier versions of this manuscript

